# In-vitro Knockdown and Pharmacological Inhibition of RIPK2 Attenuates Calcium Oxalate–Nephrocalcinosis Associated Inflammation and Oxidative Stress in NRK-52E and Primary Renal Epithelial Cells

**DOI:** 10.1101/2025.07.13.664563

**Authors:** Ganesh Panditrao Lahane, Arti Dhar, Audesh Bhat

## Abstract

Ectopic deposition of calcium oxalate in the parenchyma (nephrocalcinosis) or as stones in the collecting system (nephrolithiasis) causes inflammation and oxidative stress in renal tissue. Receptor interacting serine/threonine kinase 2 (RIPK2) is a well-known mediator of oxidative stress and inflammation. However, its role in nephrocalcinosis or nephrolithiasis remains unexplored. NRK-52E and primary renal cells were treated with calcium oxalate-monohydrate (COM) to induce nephrocalcinosis like pathological changes. siRNAs and commercially available inhibitor was used to block RIPK2’s activity. Oxidative stress, inflammation, apoptosis, crystal adhesion, and changes in cell morphology were measured as endpoint markers. Exposure to COM but not to adenine, H_2_O_2_, and high-glucose significantly upregulated RIPK2 levels and induced oxidative stress, intracellular-calcium overload, and mitochondrial dysfunction. This was accompanied by the activation of NF-κB pathway, high levels of pro-inflammatory and low levels of anti-inflammatory cytokines, increased apoptosis, and epithelial-mesenchymal transition (EMT). RIPK2 silencing or pharmacological inhibition effectively mitigated these pathological changes and restored levels of antioxidant enzymes. Mechanistically, RIPK2 inhibition disrupted the NF-κB/TGF-β1 signaling and reduced CaOx crystal adhesion. Preliminary data from our previously conducted *in vivo* CaOx mouse model study confirmed the CaOx-induced RIPK2 upregulation. Our data strongly supports the involvement of RIPK2 in CaOx-induced renal cell damage.

## 1. Introduction

Nephrocalcinosis and nephrolithiasis are two clinically distinct conditions, but they share important features, such as renal calcium deposition, oxidative stress, inflammation, and cellular injury [1, 2]. In nephrocalcinosis, calcium oxalate (CaOx), also referred as oxalosis, and calcium phosphate (CaP) crystals are deposited within the renal parenchyma. In contrast, nephrolithiasis refers to the formation of kidney stones, which are predominantly composed of CaOx and are found in the collecting systems of the kidneys [2]. Despite advancements in diagnosis and treatment, both nephrocalcinosis and nephrolithiasis remains a significant health problem due to their complex etiology, high recurrence rate, and association with chronic diseases like hypertension, diabetes mellitus, and renal failure [3]. A key factor in CaOx crystal deposition and stone formation process is the interaction between crystals and renal cells, particularly the adhesion and internalization of CaOx crystals by tubular epithelial cells. These interactions not only facilitate the retention of crystals but also trigger cellular damage and a microenvironment that promotes stone formation [4, 5].

Exposure to CaOx and crystal deposition can disrupt cellular homeostasis by promoting inflammation, mitochondrial dysfunction, and excessive production of reactive oxygen species (ROS) [6–9]. ROS-mediated oxidative stress, often regulated by NADPH oxidase (NOX) activity, impairs cell viability and activates both apoptotic and necrotic cell death pathways [10]. Additionally, oxidative stress and inflammation contribute to renal damage and play a role in the epithelial-mesenchymal transition (EMT) [11]. There is growing evidence linking kidney nephrocalcinosis/nephrolithiasis induced injury to the progression of EMT. Notably, CaOx crystals have been shown to initiate the TGF-β/Smad signaling pathway, leading to the activation of transcription factors associated with EMT [12, 13]. A pioneering study by Boonla et al. [14] first confirmed the onset of EMT in the renal tissue of patients with nephrolithiasis. Recent research has indicated that CXCR4 signaling helps to prevent CaOx-induced EMT by inhibiting the NF-κB pathway [15]. Furthermore, in a recent study, restoring SIRT6 levels in mice led to the attenuation of CaOx-induced renal inflammation and oxidative stress [16]. These findings underscore the importance of understanding the molecular mechanisms behind CaOx-induced injury and its relationship with metabolic imbalances and EMT in the pathophysiology of nephrocalcinosis/nephrolithiasis.

The Receptor-Interacting Protein Kinase (RIPK) family contains serine/threonine kinases, with RIPK1–RIPK7 as its known members [17]. RIPK2, also known as RIP2, RICK, CARDIAK, and CARD3 is an intracellular 61 kDa kinase protein containing an N-terminal kinase domain and a C-terminal caspase activation and recruitment domain (CARD) [18]. Functionally, RIPK2 is a central adaptor molecule in innate immune signaling. It transduces downstream signaling of pattern recognition receptors (PRRs), namely the nucleotide-binding oligomerization domain-containing receptors NOD1 and NOD2 [19]. Upon NOD1/2 activation, RIPK2 promotes recruitment of signaling complexes through its CARD domain, resulting in the activation of nuclear factor kappa B (NF-κB) and mitogen-activated protein kinases (MAPKs) [20, 21]. By interacting with IKKγ (NEMO), RIPK2 enhances IKK complex activation and eventual transcription of pro-inflammatory genes [22]. Importantly, RIPK2 expression is inducibly upregulated by NF-κB signaling, implying a feedback amplification loop [22, 23].

Not only in inflammation, RIPK2 also plays a crucial role in apoptosis by interacting with caspase-1 and contributes to cell death during episodes of oxygen-glucose deprivation [24]. RIPK2 is also positioned upstream of transforming growth factor-beta-activated kinase 1 (TAK1), which is a key regulator of MAPK-mediated inflammatory signaling. Both clinical and experimental evidence links RIPK2 to a variety of inflammatory and autoimmune disorders, including allergic airway inflammation [25], inflammatory bowel disease [26], and multiple sclerosis [27]. Notably, RIPK2-deficient mice (Ripk2−/−) show resistance to experimental autoimmune encephalomyelitis [27], highlighting the critical role of RIPK2 in neuroinflammation.

Recent evidence suggests that RIPK2 is involved in inflammatory and apoptotic signaling pathways across various diseases. However, its role in renal disorders has not yet been studied. To date, no research has focused on RIPK2’s involvement in COM-induced renal tubular injury or its potential link to EMT, which is a key factor in the development of fibrosis in chronic kidney disease (CKD). Given RIPK2’s established role in regulating NOD-like receptor (NLR) signaling and its reaction to oxidative stress—both of which are significant in crystal-induced nephrotoxicity—we hypothesized that RIPK2 could exacerbate COM-induced cell damage. Therefore, this study aimed to investigate the mechanistic role of RIPK2 in COM-induced renal epithelial injury.

## 2. Material and methods

### 2.1. Chemicals and reagents

Predesigned siRNA sequences targeting RIPK2/RICK (siRICK#1, siRICK#2) and a universal scrambled siRNA control (siCtrl), along with hematoxylin, eosin, Sirius Red, 2′,7′-dichlorofluorescin diacetate (DCFDA), RIPA buffer, Griess reagent, and all primers, were purchased from Sigma-Aldrich (St. Louis, Missouri, USA). Antibodies against TNF-α, p-NF-κB, and TGF-β were obtained from Novus Biologicals, LLC (Colorado, USA). Lipofectamine™ 2000 reagent, TMRE, Flura-2 AM, and FBS were purchased from Invitrogen, Thermo Fisher Scientific (Massachusetts, USA). Antibodies against RIPK2/RICK, IL-1β, Caspase-3, Vimentin, E-cadherin, GAPDH, HRP-conjugated secondary antibodies, and Alexa Fluor-conjugated antibodies were obtained from Santa Cruz Biotechnology (California, USA). Antibodies for IL-6 and IL-10 were purchased from Abclonal (Massachusetts, USA). DMEM, BSA, trypsin, MTT reagent, and DAPI were procured from HiMedia (Mumbai, India). The BCA protein assay kit, TB Green® Premix Ex Taq™ II, PrimeScript™ RT reagent kit, and RNAiso Plus were obtained from Takara Bio Inc. (Shiga, Japan). The TACS® Annexin V-FITC Apoptosis Detection Kit was purchased from R&D Systems (Minneapolis, USA).

### 2.2. Animals

Animal experiments were conducted in accordance to the Institutional Animal Ethics Committee (IAEC) of BITS-Pilani, Hyderabad Campus approved (Approval No: BITS-Hyd/IAEC-2022-53) and Animal Research Reporting of In Vivo Experiments (ARRIVE) guidelines. The *in vivo* data shown in this study were collected from our previously published study [28] and from an ongoing study (unpublished data). Briefly, male C57BL/6 mice (5–7 weeks old, weighing 20–25g) were procured from Vyas Lab, Hyderabad, India. Animals were maintained under controlled environmental conditions (22±2°C temperature, ∼60% relative humidity) with a 12h light/dark cycle. The study included two model groups; Adenine-induced CKD group (n=6) and glyoxylate-induced nephrocalcinosis group (n=6) along with their corresponding control groups (n=6). Before initiating the treatment, animals were acclimatized for one week under same conditions followed by random allocation to each group using a simple randomization method of random number generator in Microsoft Excel. Adenine-induced CKD group received 50mg/kg adenine in 0.5% CMC via oral gavage on alternate days for 30 days. The corresponding control group received vehicle (0.5% carboxymethyl cellulose) via oral gavage. The glyoxylate-induced nephrocalcinosis group received 100mg/kg/day intraperitoneal (i.p.) injections for 8 consecutive days. The corresponding control group received i.p. injections of 0.9% NaCl (vehicle). The number of animals per group was based on our previous experience in conducting similar studies, published literature, and ethical considerations to ensure adequate statistical power while minimizing animal use. Throughout the study, animals had free access to standard chow and water and were housed under identical conditions. Inclusion criteria included; healthy, wild type, specific pathogen-free (SPF) animals with no prior procedures or known health issues. Inclusion criteria were based on body weight, physical activity, and general health status during the acclimatization period. Exclusion criteria mainly involved any animal exhibiting signs of acute illness or distress that were unrelated to the treatment protocol. No specific exclusion criteria for animals or data points were formally established prior to the experiment. All animals assigned to the experimental group (n=6) completed the protocols as planned and were included in the analysis. At the end of the study, mice were euthanized, and kidneys were collected. The kidney tissues were immediately snap-frozen and stored at −80`D`°C for subsequent analysis, including qRT-PCR and Western blotting. Additionally, six healthy mice were used solely for the isolation of renal epithelial cells, which were then utilized in *in vitro* experiments.

### 2.3. NRK-52E culture

NRK-52E rat renal tubular epithelial cells, procured from the National Centre for Cell Science (NCCS, Pune, India), were maintained in DMEM with high glucose (4.5g/L), supplemented with 10% FBS and 1% penicillin-streptomycin antibiotic cocktail. The cells were incubated under standard culture conditions at 37°C in a humidified atmosphere with 5% CO`•`.

### 2.4. Mouse Primary renal tubular epithelial (PRTE) cell isolation and culture

PRTE cells were obtained by a modified enzymatic digestion procedure based on previously described methods [29]. In brief, the renal cortex of mice kidney was collected, minced, and digested with collagen type-2 at 37°C for 5min. Undigested pieces were removed by filtration, and enzymatic activity was stopped by adding 10mL of complete DMEM/F-12 growth medium to the suspension. The homogenate was subjected to sequential centrifugation (50×g, 5min) to retrieve tubular cell-enriched pellets, and then washed twice in 1x phosphate-buffered saline (PBS) to remove residual debris. Purified PRTE cells were suspended in a fresh medium and seeded on collagen-coated culture dishes to augment cellular adhesion. The cells were maintained under similar conditions as NRK-52E cells, specifically at 37°C in a humidified incubator with 5% CO`•`.

### 2.5. Cell viability assay

Cell viability and the optimal concentration of COM were assessed using the MTT assay. NRK-52E and PRTE cells were seeded in 96-well plates at a density of 1×10^4^ cells per well and allowed to grow for 24h. Cells were treated with increasing concentrations of COM (25, 50, 100, 150, 200, and 250µg/mL) for 24h. Post-treatment, 0.5mg/mL of MTT reagent was added and the cells were incubated for another 4h. The medium was then discarded, and 100µL of DMSO was added to dissolve the formazan crystals. The absorbance was measured at 570nm using a microplate reader (Spectramax M4, Molecular Devices, LLC), and cell viability was determined based on the control group.

### 2.6. Gene knockdown assay

To evaluate the functional role of RIPK2 in COM-induced renal injury, NRK-52E cells were transfected with two independent pre-designed 50nM anti-RIPK2 siRNAs (siRICK#1, siRICK#2) or a 50nM universal scrambled siRNA control (siCtrl) using Lipofectamine following the manufacturer’s protocol. Transfected cells were allowed to recover for 24h in complete DMEM medium before checking the RIPK2 knockdown efficiency. For COM treatment experiments, transfected cells were divided into six experimental groups: (1) Control: Untreated cells, (2) siRNA control: Scrambled siRNA-transfected cells (siCtrl), (3) siCtrl+COM: Scrambled siRNA-transfected cells+COM (100µg/mL, 24h), (4) COM: Untreated cells+COM (100µg/mL, 24h), (5) siRICK#1 + COM: RICK siRNA#1-transfected cells+COM (100µg/mL, 24h), and (6) siRICK#2+COM: RICK siRNA#2-transfected cells+COM (100µg/mL, 24h). Following 24h of siRNA transfection, cells in groups 3–6 were exposed to COM crystals (100µg/mL) for an additional 24h under standard culture conditions. Post-treatment, cells were harvested for downstream analysis.

### 2.7. Pharmacological inhibition of RIPK2

NRK-52E and PRTE cells were seeded into collagen-coated 6-well plates at a density of 1.5 × 10`•` cells/well and allowed to adhere for 24h under standard culture conditions. To evaluate the therapeutic potential of RIPK2 inhibition, cells were divided into four experimental groups: (1) Control: Untreated cells maintained in basal medium. (2) COM: Cells exposed to COM (100µg/mL) for 24h. and maintained in basal medium for an additional 24h. (3) COM + OD36: Cells treated with COM (100µg/mL) for 24h, followed by post-treatment with RIPK2 inhibitor OD36 (50nM) for an additional 24h. (4) OD36: Cells maintained in basal medium for 24h, followed by treatment with OD36 (50 nM) for the next 24h. Following treatments, cells were washed with PBS to remove residual crystals and inhibitors, and then harvested for downstream analyses.

### 2.8. Dichlorofluorescein (DCFH-DA) assay for ROS

A previously published protocol was followed to perform the experiment [30]. Briefly, cells were seeded in 6-well plates at 1.5 × 10^4^ cells/well density and treated with OD36 and RIPK2 siRNAs respectively as mentioned above in transfection and RICK inhibition protocol. At the end of the treatment scheduled, the cells were trypsinized, centrifuged, washed twice with ice-cold PBS, and incubated with DCFH-DA at a final concentration of 10μM in ice-cold PBS for 20min. After cells were washed three times with ice-cold PBS to remove the extracellular DCFH-DA, cells were resuspended in 300µl ice-cold PBS followed by the measurement DCF fluorescence intensity by FACS (BD FACS Aria III).

### 2.9. qPCR analysis

Total RNA was isolated from cells using TRIZOL reagent (RNAiso plus). Reverse transcription synthesis of cDNA was performed using the PrimeScript RT Reagent Kit (Takara, Cat.No.RR037A). qPCR was performed on a CFX Connect Real-Time PCR Detection System (Bio-rad, USA) using the published protocol [30]. β-actin was used as an internal control to normalize the mRNA levels of selected genes and values were presented as relative to the control taken from three independent replicates. Primer details are mentioned in **Suppl. Table 1.**

### 2.10. Measurement of mitochondrial membrane potential

Mitochondrial membrane potential (MMP) was assessed according to the previously published protocol [28]. Briefly, cells were treated under various conditions, as described in the gene knockdown and RIPK2 inhibition sections. After treatment, cells were trypsinized, centrifuged, and washed 3X with PBS. Subsequently, the cells were incubated with 400nM TMRE in PBS for 20min, followed by three additional washes with PBS, and samples analysis by FACS using the propidium iodide (PI) filter [28].

### 2.11. Apoptosis detection

Apoptosis was detected using Annexin V fluorescein isothiocyanate (FITC)/propidium iodide (PI) apoptosis detection kit according to the manufacturer’s instructions. After completion of the treatment schedule, cells were collected, washed twice with cold PBS, and incubated with 5μl of Annexin V-FITC for 15min, followed by incubation with propidium iodide (PI; 10 μl) for 10min in the dark at 4°C. Cells were run in a FACS machine to detect early apoptotic, late apoptotic, and necrotic cells papulations [30].

### 2.12. Intracellular calcium levels

Intracellular calcium (Ca²`D`) levels were measured using flow cytometry in the same treatment groups as mentioned in the gene knockdown and RIPK2 inhibition sections. After treatment, the cells were trypsinized, centrifuged, and washed 3X with calcium-free PBS. Next, the cells were incubated with 1 µM Fura-2AM at 37°C for 2–3h. Following the incubation, the cell suspensions were centrifuged and washed with calcium-free buffer. The final cell pellet was then resuspended and analyzed by flow cytometry. Corrections for autofluorescence were made by using Fura-2AM untreated cells. A minimum of 10,000 events were acquired per sample to determine intracellular calcium levels.

### 2.13. Immunofluorescence staining

Cells were seeded on glass coverslips placed in 12-well plates and cultured until 50–60% confluency. Treatments were then applied as mentioned earlier. Post-treatment period, cells were washed with PBS and fixed with 4% paraformaldehyde at room temperature (RT) for 15 minutes, followed by permeabilization with 0.1% Triton X-100 in PBS for 5 minutes. Cells were subsequently washed 2–3 times with PBS and blocked with 2% BSA for 1h at RT. Cells were incubated overnight at 4°C with selected primary antibodies. The next day, after three PBS washes, cells were incubated in the dark for 1h with the appropriate fluorophore-conjugated secondary antibody (Alexa Fluor 488 or Alexa Fluor 594). Nuclei were counterstained with DAPI. Finally, the coverslips were mounted on glass slides using an anti-fade mounting medium and imaged using a confocal laser scanning microscope (Leica DMi8, Germany).

### 2.14. Western blot analysis

Briefly, cellular and kidney tissue proteins were extracted and quantified using the BCA assay. Subsequently, 25μg of denatured protein per well separated in a 12% SDS-PAGE gel and transferred to a PVDF membrane. Membranes were blocked with 3% BSA for 1h followed by overnight incubation at 4°C with anti-RIPK2 (1:1000) and anti-GAPDH (1:1000) antibodies. After three washes with 1X PBST, membranes were incubated with HRP-conjugated secondary antibody (1:3000) for 2h at room temperature. Protein bands were visualized using the VILBER FUSION Solo S Western blot and chemiluminescence imaging system (France), and band intensities were quantified using ImageJ software (NIH, Bethesda, MD, USA). Relative protein expression was calculated as the ratio of the target protein band intensity to that of GAPDH.

### 2.15. Cell adhesion assay

Cells were cultured in 12-well plates until they reached approximately 80% confluence. Subsequently, the cells were treated as described in the transfection and RIPK2 inhibition assays. After treatment, the cells were washed thoroughly three times with PBS to remove any unbound COM crystals. Finally, bright-field images were captured using an inverted microscope, and the number of adherent crystals was counted in at least 10 randomly selected high-power fields per well.

### 2.16. H&E staining

After the treatment schedule, cells were fixed with ice-cold methanol for 10min, followed by a PBS wash. Subsequently, the cells were stained with Mayer’s hematoxylin for 1.5min, then briefly rinsed with acid alcohol to remove excess stain, followed by another PBS wash, which is a crucial step. Next, the cells were stained with eosin for 30sec. Finally, 3–4 quick PBS washes were given, and the cells were observed under an optical microscope.

### 2.17. Sirius red staining

At the end of the treatment, the culture medium was removed, and the cells were washed with PBS and fixed with ice-cold methanol at 4°C for 15mins. After 2–3 PBS washes, the cells were stained with 0.1% Sirius Red (Direct Red 80) solution prepared in 1% acetic acid. The plates were then incubated at room temperature for 1h. Subsequently, the wells were washed with 400μL of 0.1M HCl, followed by a PBS wash. Finally, images were captured using an optical microscope, and collagen deposition was analyzed using ImageJ software.

### 2.18. Statistical analysis

Data are presented as mean ± standard deviation (SD) and were analyzed using GraphPad Prism version 8.0 (GraphPad Software Inc., San Diego, CA, USA). The significant Differences among groups were evaluated using one-way ANOVA followed by the Bonferroni post hoc test. The p<0.05 was considered statistically significant.

## 3. Results

### 3.1. COM-induces RIPK2/RICK upregulation

Since RIPK2 is a known inducer of inflammation, and inflammation significantly contributes to renal cell damage [31], we hypothesized that RIPK2 may have a role in kidney damage. To test this hypothesis, we first measured RIPK2 levels in kidney tissues of glyoxylate-induced nephrocalcinosis mice (our ongoing independent study). Our findings revealed that RIPK2 levels were significantly higher in the glyoxylate-treated mice compared to the untreated mice (**Fig. 1A–C**). We also detected RIPK2 levels in kidney tissues of adenine-induced CKD mice from our recently completed study [28] and found no significant change in RIPK2 levels (**Fig. 1D–F**). These results suggested that RIPK2 may be specifically involved in glyoxylate-induced kidney damage. To validate these preliminary findings, we measured RIPK2 mRNA and protein levels in NRK-52E cells exposed to various substances, such as adenine, hydrogen peroxide (H`•`O`•`), high glucose (HG), and COM routinely used to induce renal cell injury in *in vitro* and *in vivo* model systems [28, 30]. Consistent with our *in vivo* findings, cells treated with COM exhibited the highest expression of RIPK2 among all treatments (**Fig.1G–J**), further supporting a specific role for RIPK2/RICK in COM-induced renal injury. Next, we replicated this observation in NRK-52E and PRTE cells by treating cells with COM at a concentration of 100µg/mL, as determined from a concentration gradient cytotoxicity curve (**Fig. 1K&P**). The cell viability at this concentration was 78% in NRK-52E and 76% in PRTE cells. As shown in **Fig. 1L–O and Fig. 1Q–T**, the levels of RIPK2 were significantly higher in COM-treated cells than in untreated cells.

**Fig. 1.**
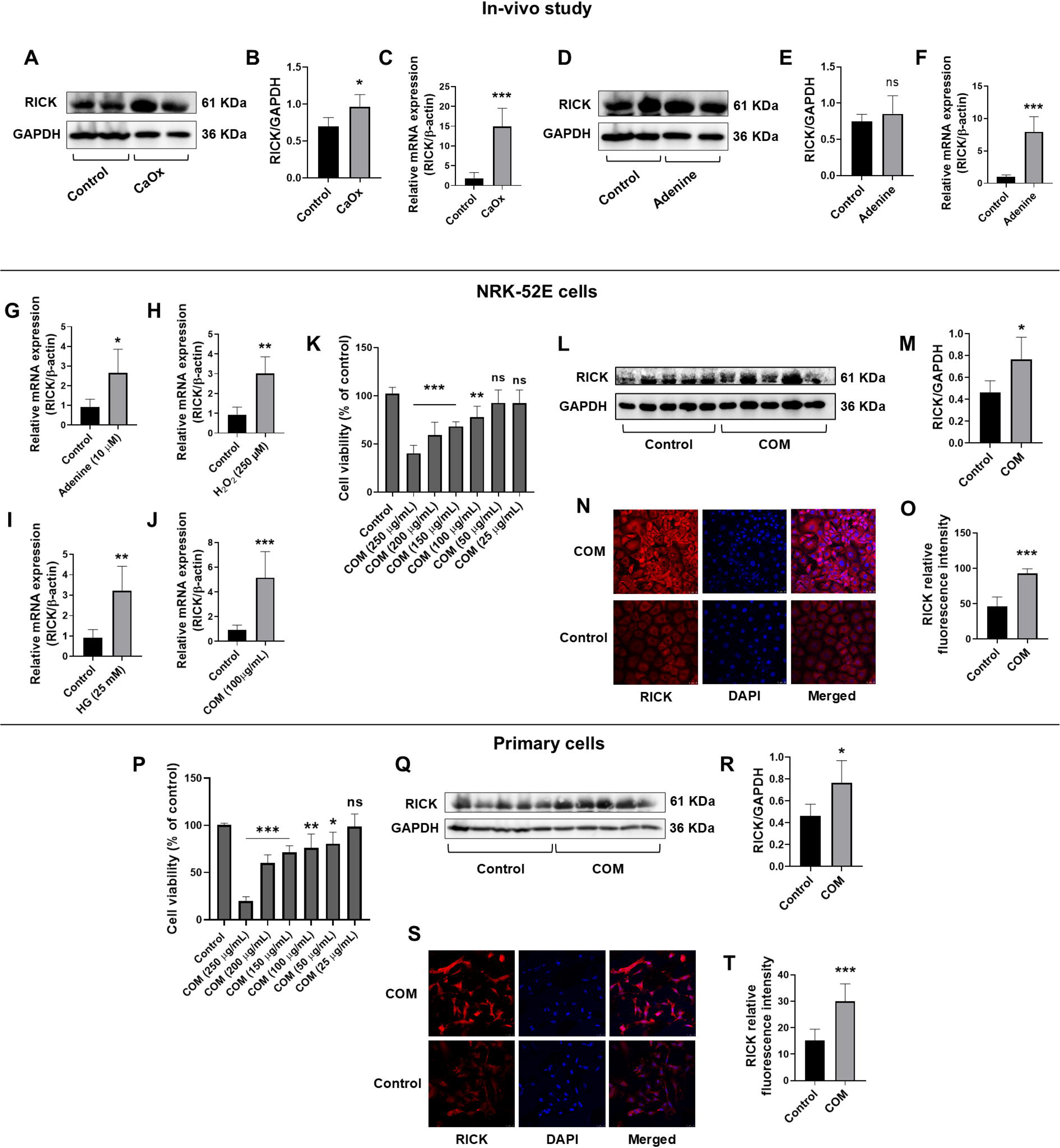
RIPK2/RICK upregulated in COM-induced NRK-52E cells and primary RTECs. (A–B) Representative western blot and quantitative analysis of RIPK2/RICK protein expression in kidneys of CaOx-induced CKD mice (n=4). (C) RIPK2 mRNA expression in kidneys of CaOx-induced CKD mice (n=5). (D–E) Representative western blot and quantitative analysis of RIPK2/RICK protein expression in kidneys of adenine-induced nephrolithiasis mice (n=2). (F) RIPK2 mRNA expression in kidneys of adenine-induced nephrolithiasis mice (n=5). (G-J) Effect of HG, H`•`O`•`, adenine, and COM on RIPK2/RICK mRNA expression in NRK-52E cells. (K and P) NRK-52E cells and primary RTECs were exposed to different concentration of COM for 24 h. then 100 µg/mL dose was selected for further studies. (L and Q) Representative western blots of RICK expression in NRK-52E cells and primary RTECs respectively. (M and R) Quantitative analysis of western blots RICK/GAPDH (n=5). (N-O and S-T) Immunofluorescence expression of RICK and relative fluorescence intensity (n=4) respective to NRK-52E cells and primary RTECs. Values are expressed as mean ± SD. Unpaired t-test was used for statistical analysis (A-J). *p < 0.05, **p < 0.01, ***p < 0.001 vs respective control groups. * p < 0.05, ** p < 0.01, and *** p < 0.001 vs respective control groups.

### 3.2. COM induces oxidative stress and increase in calcium levels via RIPK2 gene

Previous studies have shown that COM treatment induces oxidative stress and elevated intracellular calcium (Ca²`D`) levels in cell lines and kidney tissues [32, 33], we wondered if these changes are mediated through RIPK2, since our results suggest RIPK2 upregulation upon COM treatment (**Fig. 2F–H**). First, we measured the levels of ROS and intracellular Ca²`D` levels in the NRK-52E cells treated with 100 µg/mL COM and observed elevated levels of both ROS and intracellular Ca²`D`. Furthermore, COM treatment appeared to reduce the levels of catalase (CAT) as shown in **Fig. 3A–E**. To rule out that these findings were not specific to NRK-52E cells, we repeated these experiments in PRTE cells. As illustrated in **Fig. 3F–J**, we noted similar changes in ROS, intracellular Ca²`D`, and CAT levels, consistent with our findings in the NRK-52E cells. To prove that these molecular changes were indeed mediated by RIPK2, we applied two independent experimental approaches; firstly, a siRNA based RIPK2 knockdown with two different siRNAs and secondly, inhibition of RIPK2 with a commercially available inhibitor OD36 procured from Sigma Aldrich, USA. siRNA-mediated gene knockdown experiments could not be conducted in the PRTE due to poor knockdown efficiency, primarily resulting from slow cell growth. As shown in **Fig. 2A–E**, both siRNAs achieved effective RIPK2 silencing compared to the siCtrl after 24h of transfection in the NRK-52E cells. The OD36 treatment on the other hand did not cause any alteration in the RIPK2 levels in both NRK-52E and PRTE cells (**Fig. 2F–H**). When compared to the COM-alone treated cells, the RIPK2 silenced and/or inhibited COM-treated cells showed a significant reduction in levels of oxidative stress, intracellular Ca²`D` overloading, and increased CAT levels (**Fig. 3A–J**). These findings suggest the role of RIPK2 in COM-induced oxidative stress and Ca²`D` dysregulation.

**Fig. 2.**
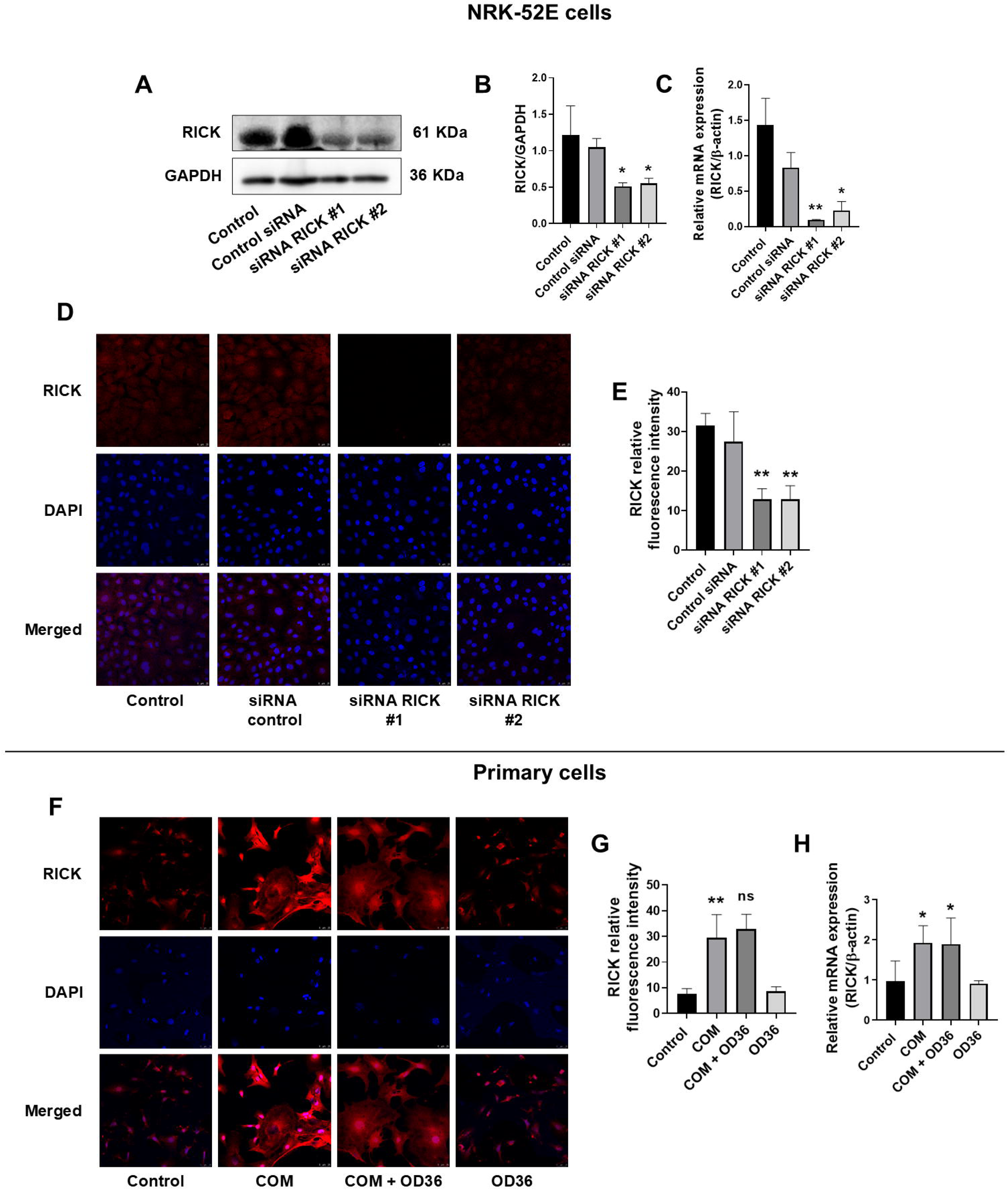
RICK Gene Silencing and pharmacological inhibition effects on mRNA and protein levels in NRK-52E cells and primary RTECs. Gene silencing effect of RICK siRNA on RICK protein and mRNA levels in NRK-52E cells were detected 24`D`h after transfection. (A-B) Representative western blots of RICK and Quantitative analysis (n=3). (C) RICK mRNA expression (n=6). (D-E) Immunofluorescence expression of RICK and relative fluorescence intensity in NRK-52E cells (n=4). (F-G) Immunofluorescence expression of RICK and relative fluorescence intensity in primary RTECs (n=4). (H) RICK mRNA expression in primary RTECs (n=6). Values are expressed as mean ± SD. * p < 0.05 and ** p < 0.01 vs respective control and siCtrl groups.

**Fig. 3.**
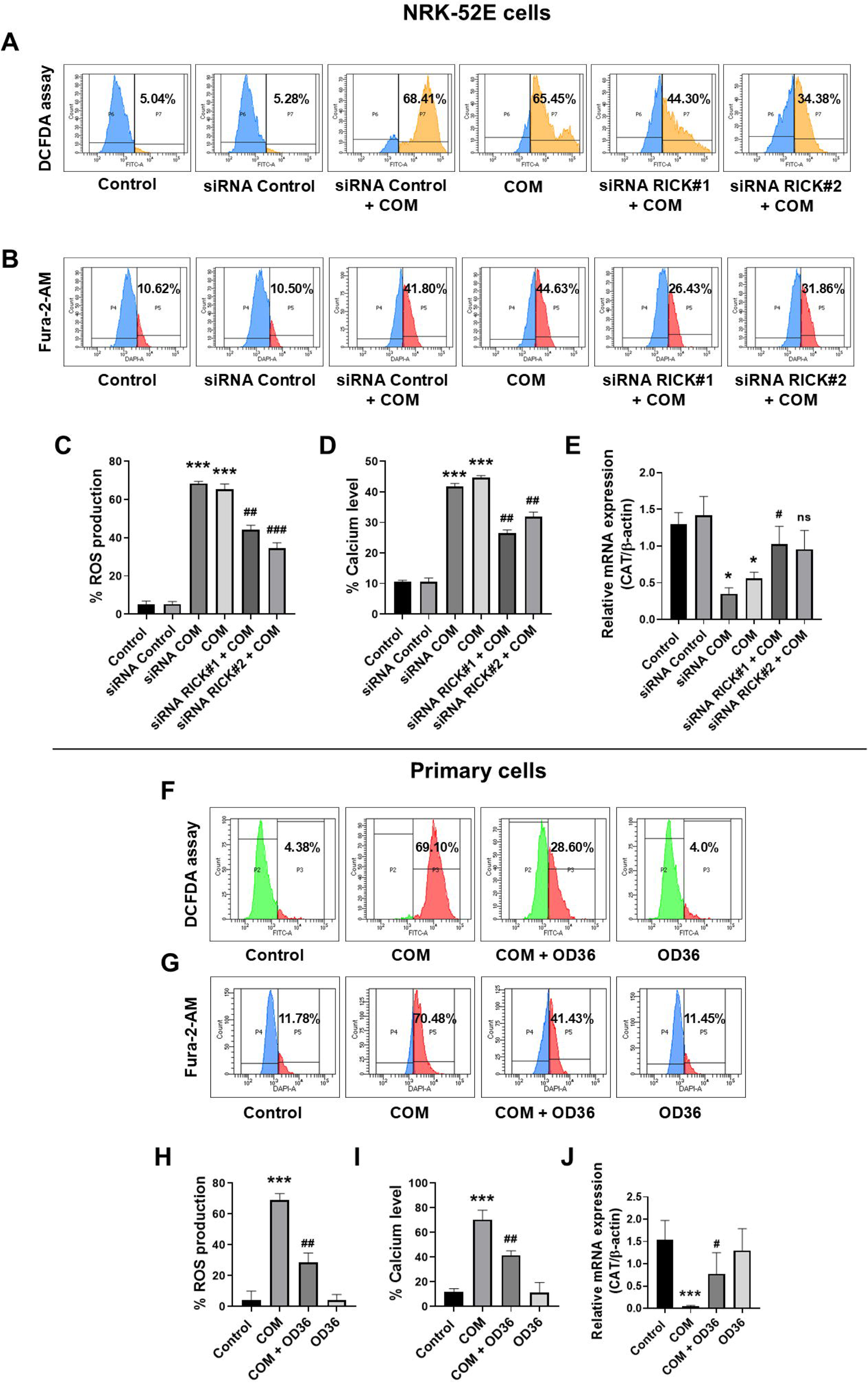
RICK Gene Silencing and pharmacological inhibition effects on COM-induced oxidative stress markers and calcium level in NRK-52E cells and primary RTECs. After 24 h of siRNA transfection, COM was exposed to NRK-52E cells for additional 24 h. (A-B) FACS-DCFDA assay and Fura-2AM assay for ROS production and Intracellular Ca^2+^ level respectively in NRK-52E cells (n=5). (C-D) Percent (%) ROS production and % intracellular Ca^2+^ level in NRK-52E cells. (E) CAT mRNA expression level in NRK-52E cells (n=6). (F-G) FACS-DCFDA assay and Fura-2AM assay for ROS production and Intracellular Ca^2+^ level respectively in primary RTECs (n=5). (H-I) % ROS production and % intracellular Ca^2+^ level in primary RTECs. (J) CAT mRNA expression level in primary RTECs (n=6). Values are expressed as mean ± SD. * p < 0.05 and *** p < 0.001 vs respective control and siCtrl groups; # p < 0.05, ## p < 0.01, and ### p < 0.001 vs respective COM groups.

### 3.3. RICK silencing and inhibition attenuates COM-induced inflammatory markers

After our initial confirmation of RIPK2’s role in oxidative stress and intracellular Ca²`D` dysregulation, we next focused on the pro-inflammatory markers, as inflammation is recognized as a key contributor to the development of nephrolithiasis. First, we investigated whether COM induces upregulation of inflammatory markers followed by the detection of RIPK2’s role using gene silencing and protein inhibition approaches. Compared to the control groups, COM treatment significantly upregulated pro-inflammatory cytokines NF-κB, TNF-α, IL-1β, and IL-6 and downregulated anti-inflammatory cytokine IL-10 in the NRK-52E cells (**Fig. 4A–K**). Similar outcomes were obtained when PRTE cells were treated with COM and OD36 (**Fig. 5A–K**). Upon RIPK2 silencing and/or inhibition, the COM-induced upregulation of NF-κB, TNF-α, IL-1β, and IL-6 and the downregulation of IL-10 was significantly attenuated (**Fig. 4 and Fig. 5**). These data further confirm that COM-induced inflammation is mediated by RIPK2.

**Fig. 4.**
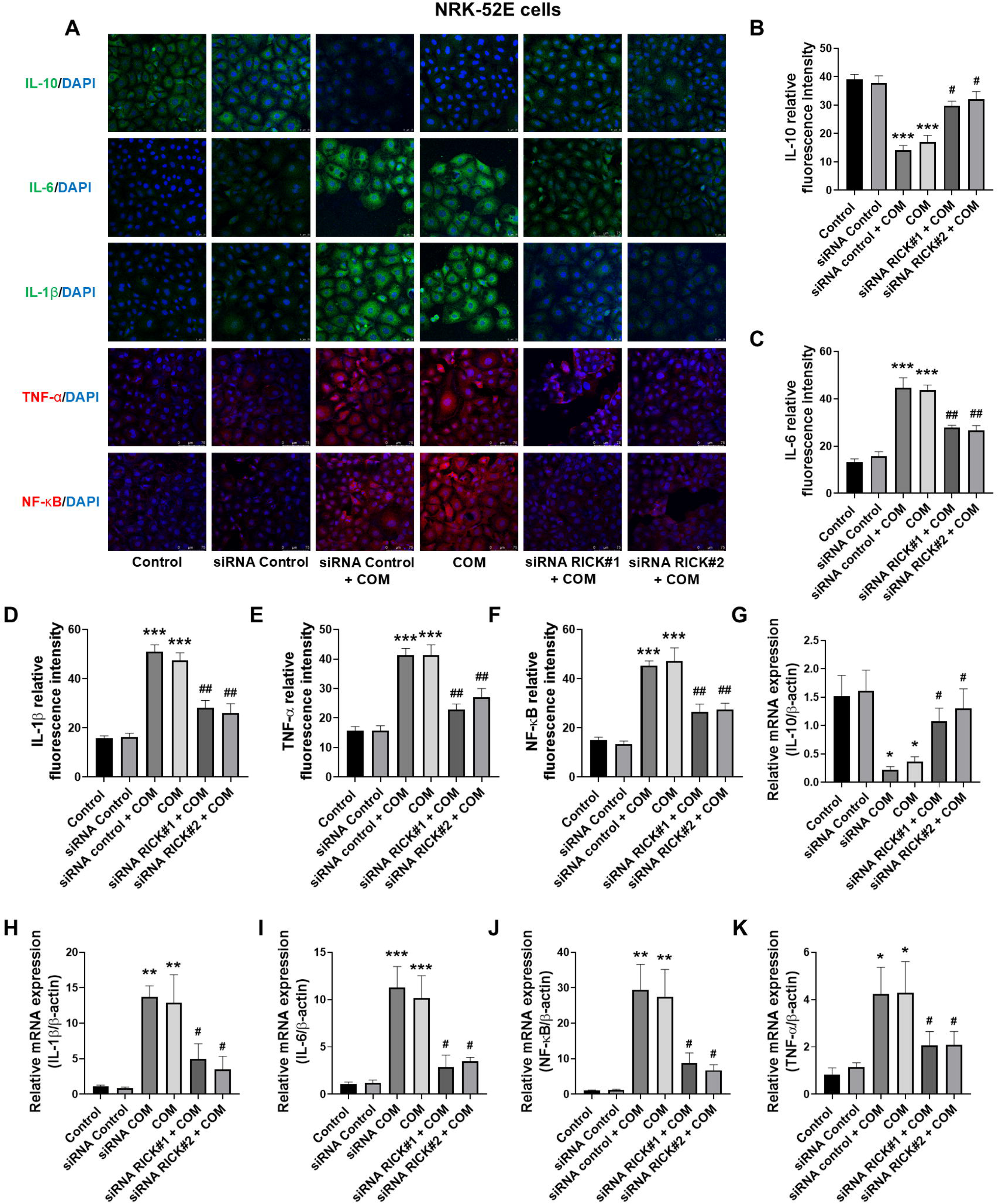
RICK Gene silencing effects on COM-induced inflammatory markers in NRK-52E cells. (A) Immunofluorescence expression of NF-κB, TNF-α, IL-1β, IL-6, and IL-10 in NRK-52E cells. (B-F) Relative fluorescence intensity of NF-κB, TNF-α, IL-1β, IL-6, and IL-10 (n=4). (G-K) The mRNA expression levels of IL-10, IL-1β, IL-6, NF-κB, and TNF-α in NRK-52E cells respectively (n=6). Values are expressed as mean ± SD. * p < 0.05, ** p < 0.01, and *** p < 0.001 vs respective control and siCtrl groups; # p < 0.05 and ## p < 0.01 vs respective COM groups.

**Fig. 5.**
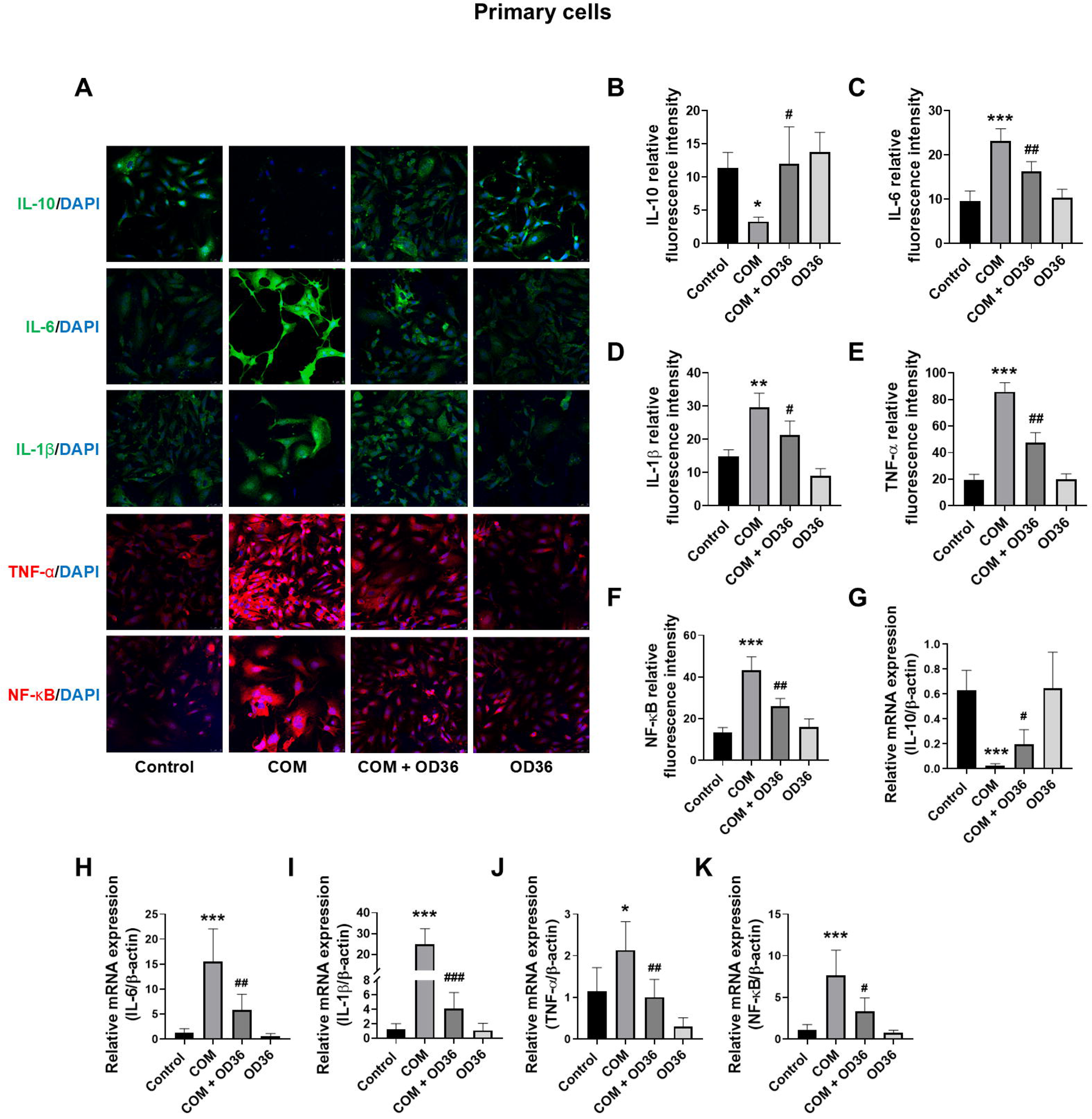
RICK inhibition effects on COM-induced inflammatory markers in primary RTECs. (A) Immunofluorescence expression of NF-κB, TNF-α, IL-1β, IL-6, and IL-10 in primary RTECs. (B-F) Relative fluorescence intensity of NF-κB, TNF-α, IL-1β, IL-6, and IL-10 (n=4). (G-K) The mRNA expression levels of IL-10, IL-6, IL-1β, TNF-α, and NF-κB in primary RTECs respectively (n=6). Values are expressed as mean ± SD. * p < 0.05 and *** p < 0.001 vs respective control groups; # p < 0.05, ## p < 0.01, and ### p < 0.001 vs respective COM groups.

### 3.4. RICK mediates COM-induced mitochondrial dysfunction and apoptosis

Mitochondrial dysfunction is an early indicator of oxidative stress-induced apoptosis; therefore, we first assessed the impact of COM on mitochondrial function and apoptosis followed by possible phenotype reversal upon RIPK2 silencing and inhibition. As can be seen in the TMRE histograms (**Fig. 6A&B and Fig. 7A&B**), a significant leftward shift in fluorescence peaks was observed in the COM-treated cells compared to the siCtrl and untreated control cells, confirming COM-mediated loss of mitochondrial membrane potential (MMP). Furthermore, the apoptotic effect of COM, as assessed by flow cytometry after cells were stained with Annexin V-FITC/PI, showed a markedly increased population of early apoptotic (Q4), late apoptotic (Q2), and necrotic cells (Q1) in comparison to siCtrl and untreated cells **(Fig. 6C&D and 7C&D)**. The knockdown and inhibition of RIPK2 in NRK-52E cells on the other hand significantly restored MMP and reduced the number of apoptotic and necrotic NRK-52E cells (**Fig. 6A–D and Fig. 7A–D**). To further confirm COM-induced cell apoptosis, we measured caspase-3 levels and detected higher caspase-3 levels in the COM treated cells than in the untreated cells (**Fig. 6E–G and Fig. 7E–G)**. Upon RIPK2 silencing and inhibition, the COM-induced caspace-3 levels were significantly down in comparison to the COM-treated cells (**Fig. 6E–G and Fig. 7E–G**). To rule out any cell-specific phenomena, all experiments were repeated in PRTE cells, except for the siRNA-mediated gene knockdown. As shown in **Fig. 7A–D**, all results were successfully replicated in the PRTE cells, thereby validating our findings from the NRK-52E cells. These findings collectively suggest that RIPK2 silencing mitigates COM-induced mitochondrial dysfunction and suppresses apoptosis in renal epithelial cells.

**Fig. 6.**
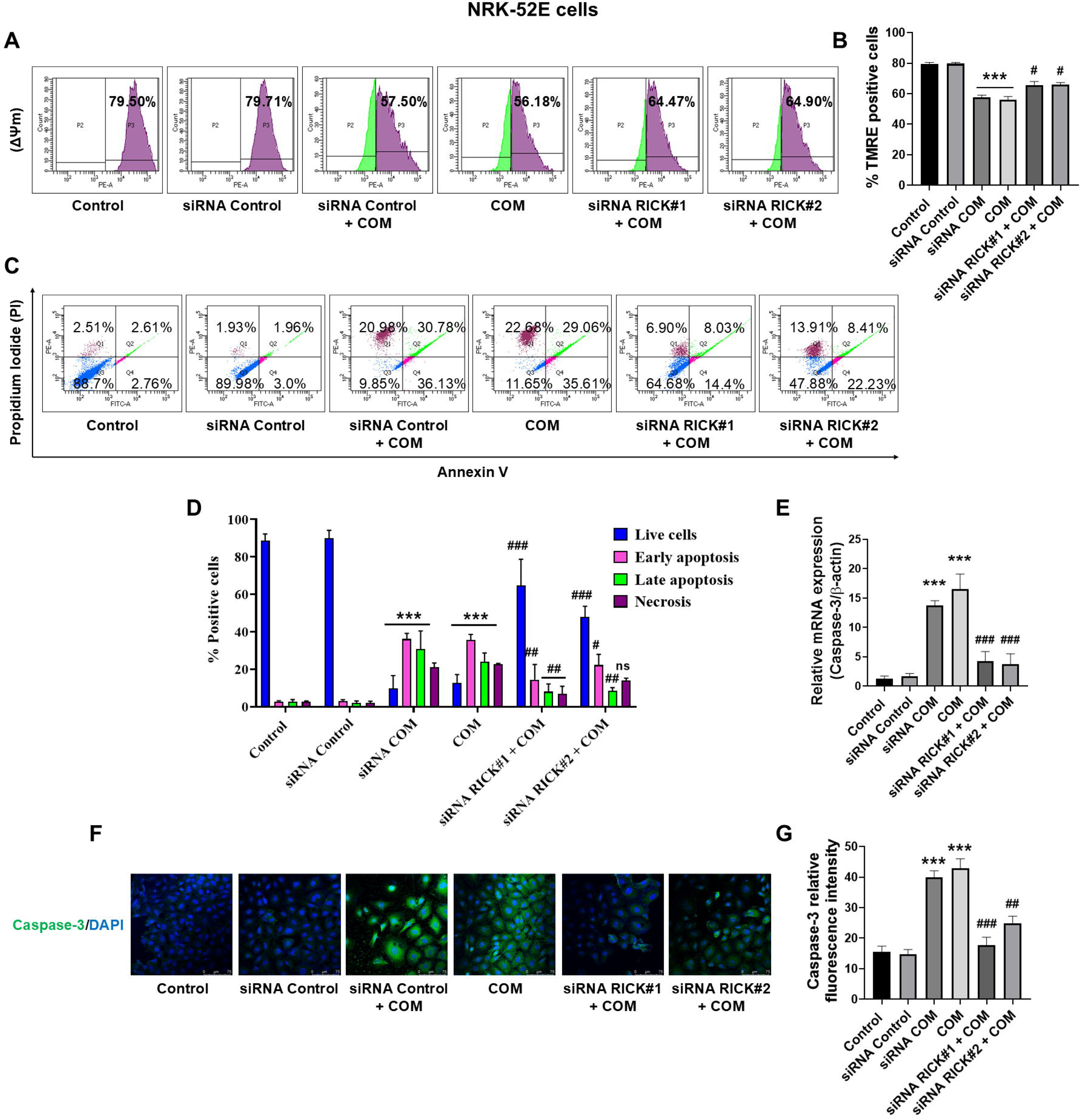
RICK Gene silencing effects on COM-induced mitochondrial dysfunction and apoptosis in NRK-52E cells. (A) TMRE dye was used for mitochondrial membrane potential by FACS in NRK-52E cells. (B) TMRE (%) positive cells (n=5). (C) The NRK-52E cells were stained with Annexin `•`-FITC/PI and analyzed by flow cytometry to determine the apoptosis population (n=5). (D) Quantitative data for the percentage of Healthy (Q3) early (Q4), late (Q2), and necrotic cells. (E) Caspase-3 mRNA expression level (n=6). (F) Immunofluorescence expression of caspase-3. (G) Relative fluorescence intensity of caspase-3 in NRK-52E cells (n=4). Values are expressed as mean ± SD. *** p < 0.001 vs respective control and siCtrl groups; # p < 0.05, ## p < 0.01, and ### p < 0.001 vs respective COM groups.

**Fig. 7.**
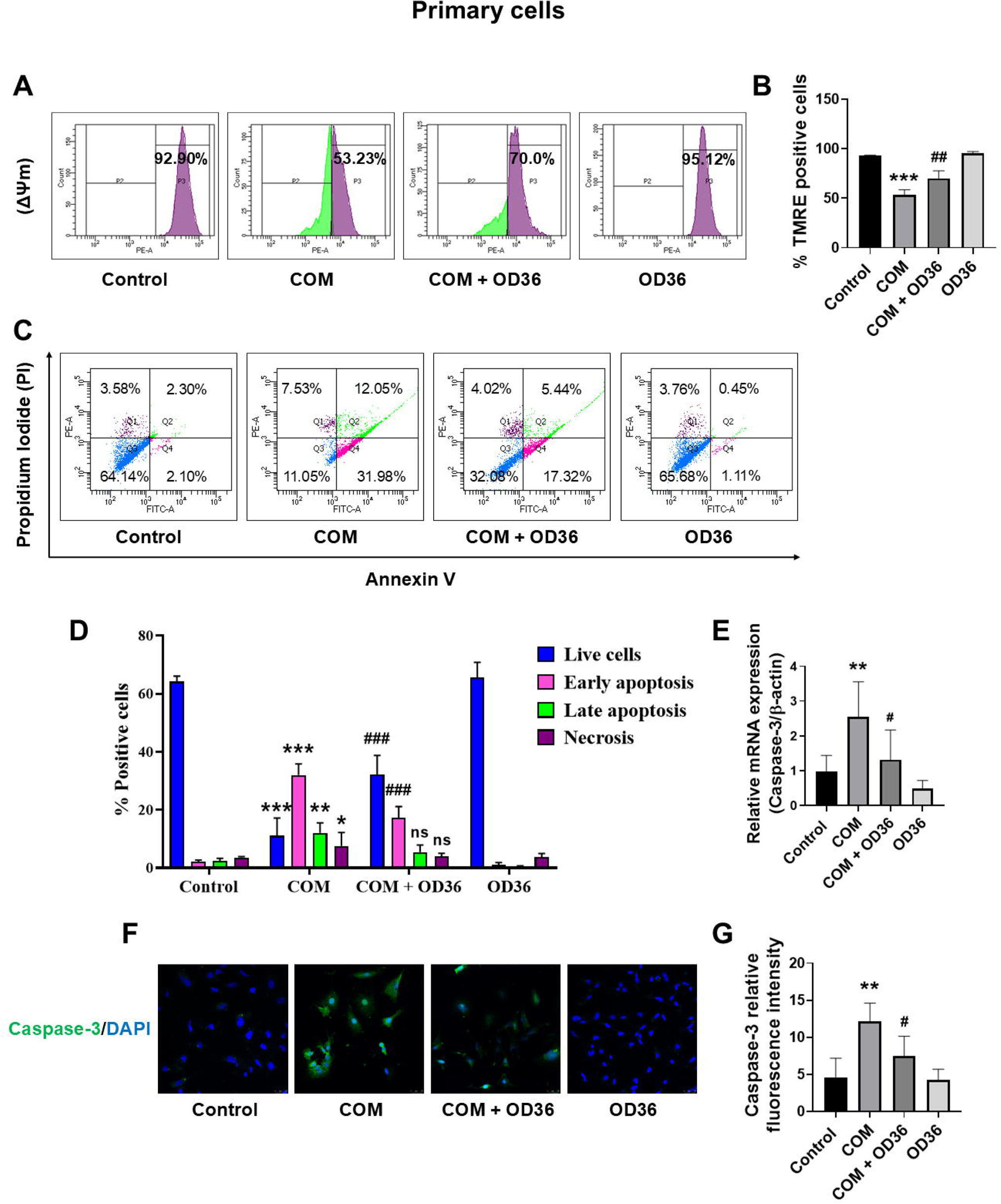
RICK inhibition effects on COM-induced mitochondrial dysfunction and apoptosis in primary RTECs. (A) TMRE dye was used for mitochondrial membrane potential by FACS in primary RTECs. (B) TMRE (%) positive cells (n=5). (C) The primary RTECs were stained with Annexin `•`-FITC/PI and analyzed by flow cytometry to determine the apoptosis population (n=5). (D) Quantitative data for the percentage of Healthy (Q3) early (Q4), late (Q2), and necrotic cells. (E) Caspase-3 mRNA expression level (n=6). (F) Immunofluorescence expression of caspase-3. (G) Relative fluorescence intensity of caspase-3 in primary RTECs (n=4). Values are expressed as mean ± SD. ** p < 0.01 and *** p < 0.001 vs respective control groups; # p < 0.05 and ## p < 0.01 vs respective COM groups.

### 3.5. RIPK2 facilitates CaOx crystal adhesion, collagen deposition, and cellular alterations

To assess the role of RIPK2 in COM-induced cellular changes in the morphology, NRK-52E cells were treated with siRNA or OD36 for 24h followed by 24h treatment with COM and PRTE cells were treated with OD36 for 24h followed by 24h exposure to COM. As revealed by the H&E and Sirius red staining, the COM-treated cells showed increased cell and nuclear size, disrupted adhesion, widened intercellular spaces, and elevated collagen deposition in comparison to the siCtrl-treated and untreated cells (**Fig. 8A–F**) and these morphological changes were significantly reversed when RIPK2 was either silenced or inhibited (**Fig. 8A–F**). Similar outcomes were observed in the PRTE cells when treated with OD36 (**Fig. 8G–L)**

**Fig. 8.**
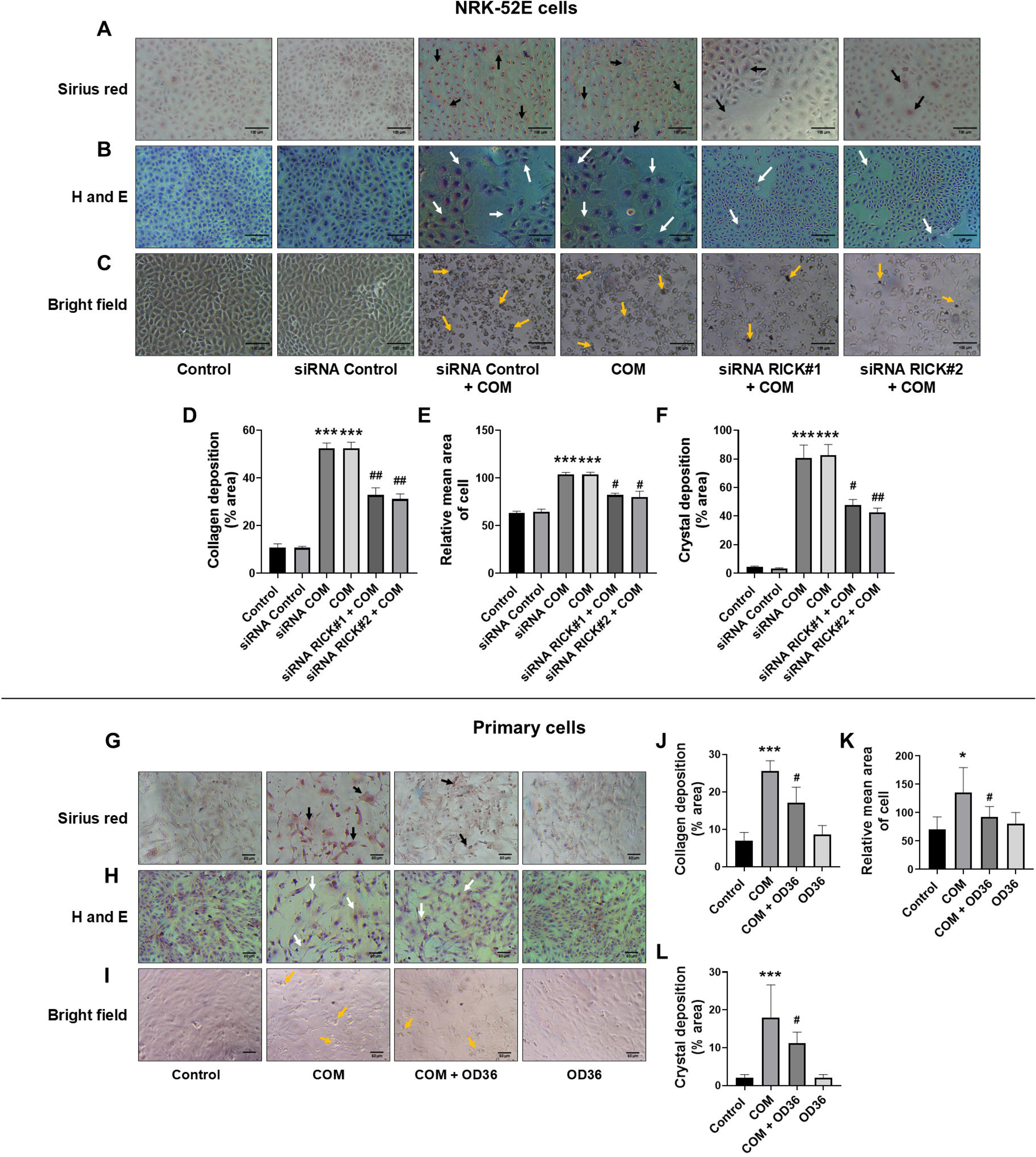
RICK gene silencing and inhibition attenuate CaOx crystal adhesion, Collagen deposition, and morphological alterations in NRK-52E cells and primary RTECs. (A and G) Sirius red staining for collagen in NRK-52E cells and primary RTECs respectively. (B and H) H and E staining in NRK-52E cells and primary RTECs respectively. Result clearly shows the decreased in cell integrity, morphology, and viability in NRK-52E cells and primary RTECs exposed COM and increased collagen secretion and deposition (Black arrow indicate the collagen and white arrow indicate the abnormal structure of cells), whereas in RICK Gene Silencing and inhibition groups reversed collagen secretion, abnormality, integrity, and nuclear condensation. (C and I) Representative bright field CaOx crystal adhesion images in NRK-52E and primary RTECs respectively (orange arrow indicate the CaOx crystals). (D and J) Collagen deposition (%) (n=4). (E and K) Relative mean area of cells (n=4). (F and L) Crystal deposition in % area (n=4). Magnification ×`D`200. Values are expressed as mean ± SD. * p < 0.05 and *** p < 0.001 vs respective control and siCtrl groups; # p < 0.05 and ## p < 0.01 vs respective COM groups.

Additionally, CaOx crystal adhesion studies showed increased crystal attachment and cell damage in COM-treated cells compared to siCtrl-treated and untreated cells. However, RIPK2 silencing or inhibition resulted in markedly reduced crystal adherence and preservation of cell viability and morphology (**Fig. 8C,F,I&L**). These findings suggest that RIPK2 modulates COM-induced morphological damage, collagen deposition, and crystal interaction in renal epithelial cells.

### 3.6. RIPK2 participates in the CaOx-induced fibrosis and EMT

As renal cells of patients with nephrocalcinosis often show signs of fibrosis and EMT, we next evaluated the role of RIPK2 in COM-induced renal fibrosis and EMT by assessing the key markers of fibroblast activation, interstitial fibrosis, and collagen deposition. Compared to siCtrl and untreated control cells, COM-exposed NRK-52E and PRTE cells exhibited high levels of TGF-β1 and vimentin staining, along with a marked reduction in E-cadherin expression (**Fig. 9A–D and Fig. 10A–D**). Correspondingly, mRNA levels of TGF-β, vimentin, collagen IV, and α-SMA were significantly upregulated, while E-cadherin mRNA level was significantly downregulated in the COM-treated cells (**Fig. 9E–I and Fig. 10E–I**). However, when RIPK2 depleted and/or inhibited cells were challenged with COM, there observed a significant reduction in the levels of fibrosis and EMT markers (**Figs. 9&10**). These data suggest that RIPK2 protein regulates COM-induced alterations in the renal cell morphology.

**Fig. 9.**
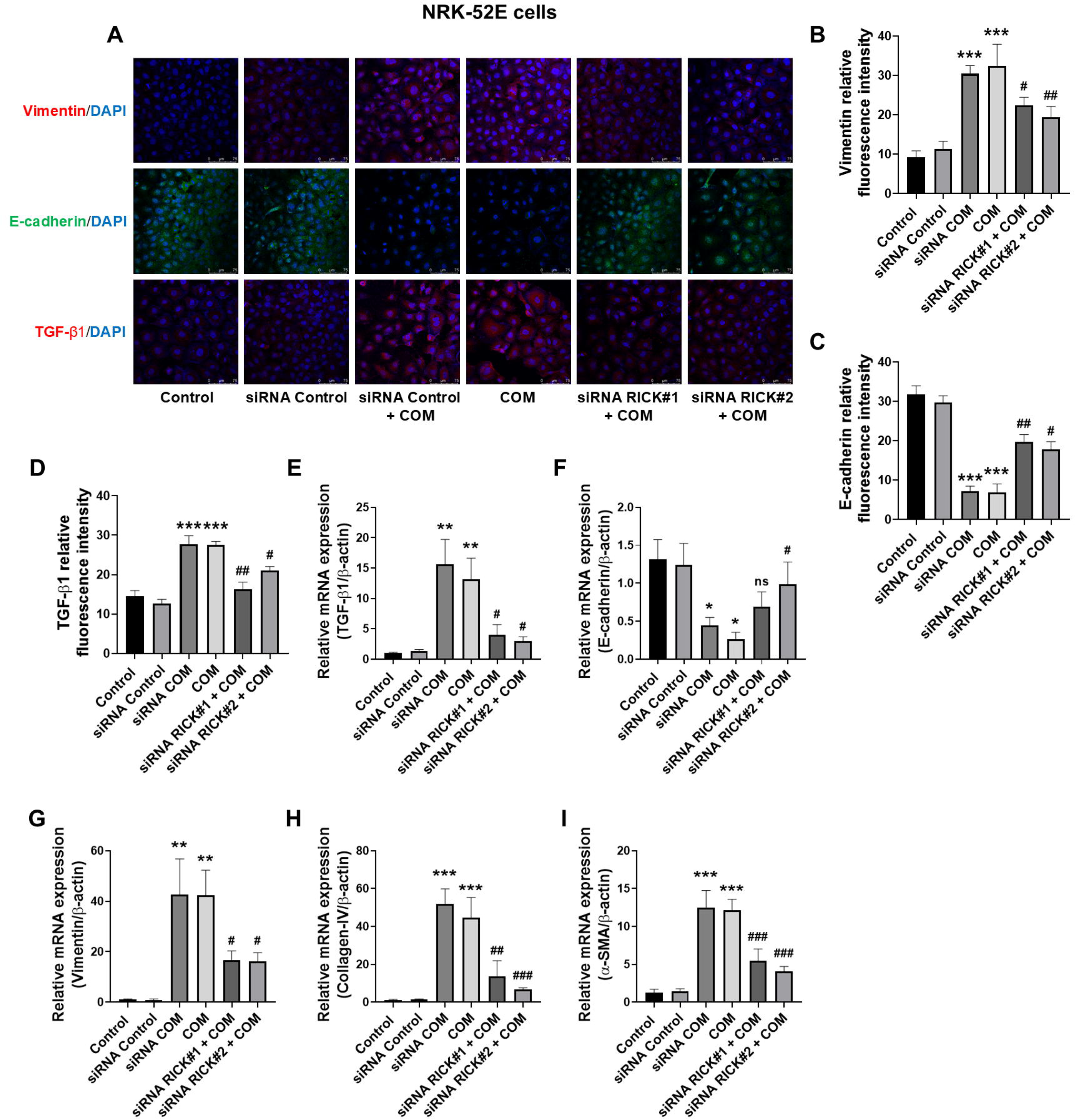
RICK Gene silencing effects on COM-induced fibrosis and EMT in NRK-52E cells. (A) Immunofluorescence expression of TGF-β, E-cadherin, and Vimentin in NRK-52E cells. (B-D) Relative fluorescence intensity of TGF-β, E-cadherin, and Vimentin (n=4). (E-I) The mRNA expression levels of TGF-β, E-cadherin, Vimentin, Collagen-IV, and α-SMA in NRK-52E cells respectively (n=6). Values are expressed as mean ± SD. * p < 0.05, ** p < 0.01, and *** p < 0.001 vs respective control and siCtrl groups; # p < 0.05, ## p < 0.01, ### p < 0.001 vs respective COM groups.

**Fig. 10.**
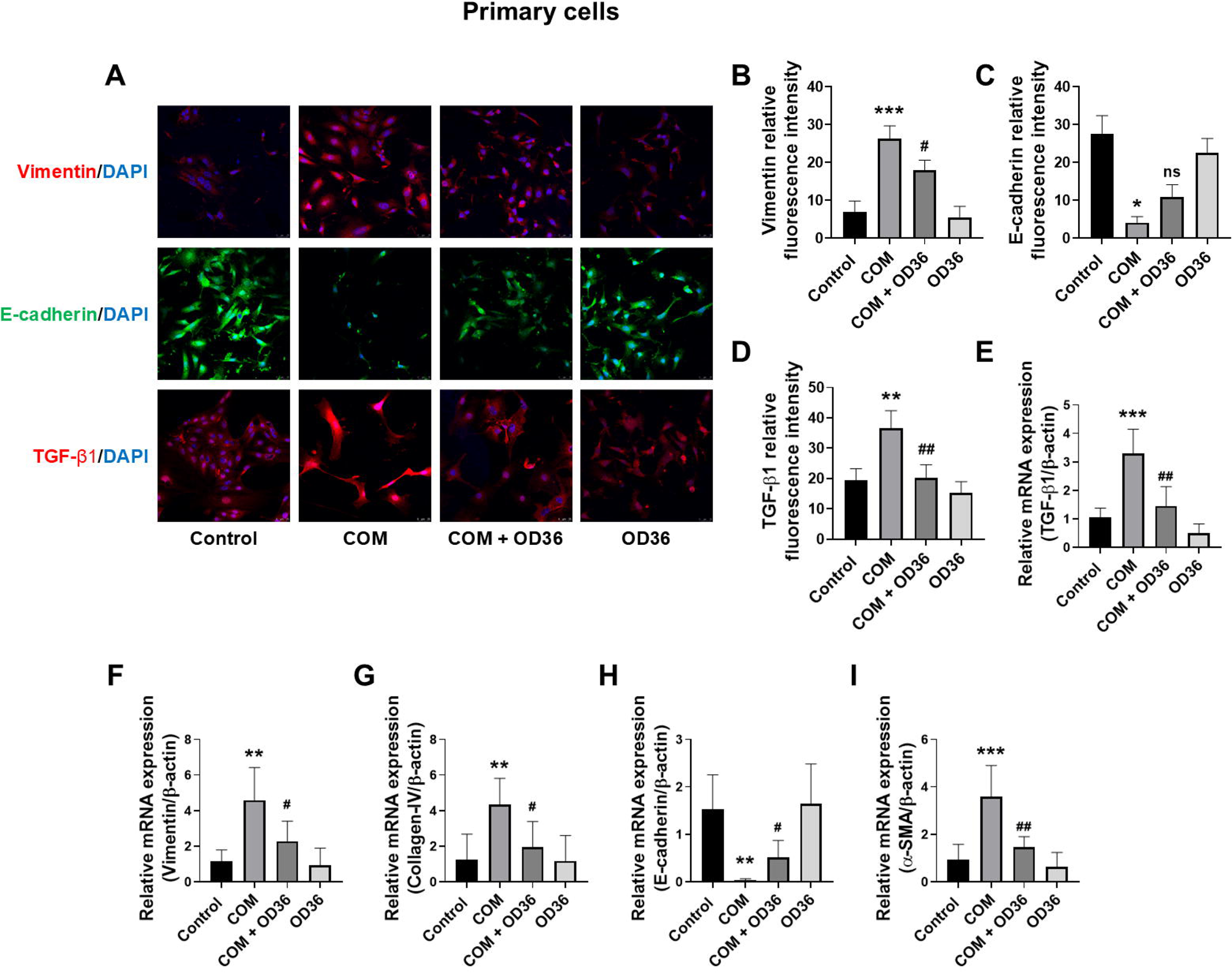
RICK inhibition effects on COM-induced fibrosis and EMT in primary RTECs. (A) Immunofluorescence expression of TGF-β, E-cadherin, and Vimentin in NRK-52E cells. (B-D) Relative fluorescence intensity of TGF-β, E-cadherin, and Vimentin (n=4). (E-I) The mRNA expression levels of TGF-β, Vimentin, Collagen-IV, E-cadherin, and α-SMA in primary RTECs respectively (n=6). Values are expressed as mean ± SD. * p < 0.05, ** p < 0.01, and *** p < 0.001 vs respective control groups; # p < 0.05 and ## p < 0.01 vs respective COM groups.

## 4. Discussion

Nephrolithiasis continues to pose a significant global health burden, affecting up to 20% of the adult population [34]. Among the various types of kidney stones, CaOx stones are by far the most prevalent, reflecting their complex, multifactorial etiology and the challenges they pose for effective prevention [35]. CaOx crystal deposition not only underlies kidney stone formation but also induces renal tubular injury, inflammation, fibrosis and EMT, all contributors of nephrocalcinosis/nephrolithiasis and progressive renal failure [36]. Although surgical and endoscopic methods have advanced stone removal procedure, effective pharmacological intervention to protect against CaOx`D`induced tubular damage and fibrotic remodeling are still lacking [35].

RIPK2 has been shown to be associated with the progression of various inflammatory and immune-related diseases and has emerged as a promising target for cancer therapy [22, 23],[25]. However, its biological role in kidney diseases, particularly nephrocalcinosis/nephrolithiasis, remains unexplored. To answer this question, our initial experimental approach of detecting RIPK2 levels in the *in vivo* kidney disease models followed by its validation in the *in vitro* model of NRK-52E cells treated with HG, H`•`O`•`, adenine, and COM-agents commonly used to induce kidney damage [28, 30, 37], provided enough evidence that perhaps CaOx stone formation and the associated renal cell damage involves RIPK2 mediated pathological changes (**Fig. 1**). For our subsequent aim to delineate the potential molecular mechanism(s) involving RIPK2, we used two different *in vitro* cell systems; a transformed normal rat kidney NRK-52E cells and an in-house generated normal mouse primary renal tubular epithelial cells to replicate our findings. Additionally, we employed two independent siRNAs to deplete RIPK2 and a commercially available RIPK2 inhibitor to replicate our results. In addition to upregulating RIPK2, COM treated cells showed signs of inflammation, apoptosis, fibrosis and EMT, all important hallmarks of nephrocalcinosis and nephrolithiasis. The question of whether these phenotypes were truly dependent on RIPK2 was addressed when we observed a significant reversal of COM-induced changes following both genetic silencing and pharmacological inhibition of RIPK2. These findings highlight RIPK2 as a potential mediator of COM-induced nephrocalcinosis-associated phenotypes and as a potential target for therapeutic intervention in CaOx-driven kidney injury. Studies have shown that inflammation and oxidative stress are closely associated with the formation of kidney stones [6, 38]. Oxidative stress primarily results from excessive production of free radicals and the release of ROS from the mitochondria [39]. Under pathological conditions, mitochondrial function is impaired due to overproduction of ROS, leading to mitochondrial DNA damage and oxidative modification of mitochondrial proteins and enzymes involved in the ATP synthesis pathway [40]. Superoxide dismutase (SOD) plays a key role in catalyzing the dismutation of superoxide anions (O_2_·^−^) into H_2_O_2_ and molecular oxygen (O_2_) [41]. During abnormal oxidative stress, SOD helps protect cells from oxidative damage by scavenging ROS [41]. CAT provides a secondary defence by converting H_2_O_2_ into water and molecular oxygen [42]. Our data clearly supports role of RIPK2 in COM-induced oxidative stress, as COM-exposure alone increased ROS production and decreased CAT production but when cells pre-treated either with siRNAs or OD36 were challenged with COM, these alterations were effectively reversed (**Fig. 3**). These findings are in concordance with the published data, as studies have shown that RIPK2 modulates oxidative stress markers and cigarette smoke-induced lung inflammation [43] and ROS levels in both myocardial ischemia-reperfusion (MI/R) injury in rats and oxygen-glucose deprivation/reperfusion (OGD/R)-stimulated cardiomyocytes [44].

Intracellular Ca²`D` is a critical second messenger involved in regulating various signaling pathways under both physiological and pathological conditions [45]. The Ca²`D` signaling plays an essential role in ROS production, as elevated intracellular Ca²`D` levels can activate ROS-generating enzymes, thereby contributing to oxidative stress [46]. The rise in cytoplasmic Ca²`D` levels is primarily mediated by Ca²`D` influx from the extracellular space and release from internal stores [45]. CaOx crystals can directly interact with and damage renal tubular epithelial cells, initiating a cascade of cellular events including increased intracellular Ca²`D` levels, oxidative stress, and cell death [36]. The high levels of intracellular Ca²`D` we observed in the COM-treated cells, followed by its attenuation upon RIPK2 silencing and inhibition suggests that RIPK2 is involved in the COM-induced dysregulation of calcium homeostasis in the kidney cells (**Fig. 3B&G**). The COM-induced upregulation of intracellular Ca²`D` is consistent with a recently published study showing increased intracellular Ca²`D` levels in NRK-52E cells following exposure to CaOx crystals [33].

Furthermore, The NF-κB signaling pathway plays a crucial role in regulating cell growth and survival, stress responses, and inflammation [47]. In addition to oxidative stress and hyperoxaluria, CaOx crystals can induce inflammatory response by activating the pyrin domain-containing NLR family member 3 (NLRP3) inflammasome, leading to the release of pro-inflammatory cytokines such as IL-1β and IL-18 [48]. Studies have consistently demonstrated the simultaneous presence of oxidative stress and inflammation in the pathogenesis of nephrocalcinosis and nephrolithiasis [6]. Excessive production of ROS can activate the NF-κB inflammatory signaling pathway and promote cellular apoptosis. Previous research has shown that inhibition of ROS production mitigates COM-induced inflammation in renal tubular epithelial cells [49, 50]. The RIPK2 has emerged as a critical regulator of inflammatory signaling pathways and has gained significant attention in recent times [51]. Elevated RIPK2 expression has been reported in patients suffering from inflammatory bowel disease [52] and genetic deletion of RIPK2 has been shown to enhance resistance to experimental autoimmune encephalomyelitis in mice [27]. Additionally, RIPK2 may function as a pro-inflammatory gene, as its knockdown has been found to protect enterocytes from colitis-induced damage [53]. To further delineate the potential link between RIPK2 and nephrocalcinosis, our investigation of inflammatory markers revealed that COM induces the expression of pro-inflammatory cytokines, such as NF-κB, TNF-α, IL-1β, and IL-6, and blocks the expression of anti-inflammatory cytokine IL-10 and when RIPK2 was either silenced or inhibited, these markers reverted significantly even when COM was present. (**Fig. 4&5**). This is the first evidence showing that COM-induced inflammatory response is mediated through RIPK2 in the renal cells. Supporting our findings, a previous study showed that intratracheal administration of RIPK2 siRNA in the lungs inhibits cigarette smoke-induced NF-κB nuclear accumulation and suppresses the expression of NF-κB-dependent pro-inflammatory mediators in the airways [43].

An increase in CaOx deposition and the associated rise in ROS weakens mitochondrial integrity, decreases MMP, and increases mitochondrial membrane permeability. Reduction in MMP activates caspases, leading to apoptosis [54]. Previous studies have also established a link between CaOx crystals, oxidative stress, DNA damage, and renal cell apoptosis [30, 39, 55, 56]. Our data suggests that COM decreases MMP and causes cell apoptosis and that these pathological changes are significantly reversed when RIPK2 is either silenced or inhibited. These data strongly support the notion that COM-induced MMP changes and cell apoptosis is dependent on RIPK2. The increased levels of caspase-3 in COM-treated cells reduced significantly when RIPK2 was either silenced or inhibited, thus confirming the role of RIPK2 in COM-induced cell apoptosis (**Fig. 6&7**). These findings are consistent with a previous report showing that RIPK2 inhibition reduces the number of TUNEL-positive cells and cleaved caspase-3 expression in *in vitro* and *in vivo* models of MI/R injury [44].

Kidney stones originate from the localized accumulation of crystals. As these crystals grow in size and interact with surrounding tissue structures, they may chemically bind to the tissues, forming adherent crystal aggregates [57]. Adhesion is a critical step in stone formation, as it not only initiates the development of kidney stones but also promotes their continued growth [4, 57]. When crystals adhere to the epithelial cells lining the renal tubules, they act as nucleation sites, encouraging further crystal deposition and thereby, increasing stone size over time [4, 57]. These cell-crystal interactions have been shown to trigger renal inflammation, leading to oxidative stress and activation of inflammatory cascades, which in turn facilitate calcium crystal aggregation and growth, ultimately contributing to kidney injury [58]. Moreover, cell injury induced by COM crystals further enhance crystal adhesion, creating a vicious cycle that accelerates kidney stone formation [58]. Our data on increased CaOx crystal adhesion to NRK-52E and PRTE cells leading to a significant damage to renal cells is in line with the published data; however, the abrogation of CaOx crystal adhesion and lack of renal cell damage in RIPK2 silenced or inhibited cells suggests that RIPK2 may play a significant role in CaOx crystal adhesion induced renal cell damage (**Fig. 8**). Moreover, kidney stone formation and nephrocalcinosis are associated with increased collagen secretion by renal tubular epithelial cells, primarily due to the damaging effects of CaOx crystals and other contributing factors [59]. In addition to CaOx crystals and oxalate, oxidative stress and inflammation also play a significant role in promoting collagen production in renal tubular epithelial cells [6, 59]. In our study, we observed elevated collagen secretion and deposition in NRK-52E and PRTE cells upon exposure to COM and that these phenotypes were effectively abrogated upon RIPK2 inhibition further support RIPK2’s role in renal cell damage (**Fig. 8**).

High oxalate conditions and COM crystals are well-established inducers of apoptosis, oxidative stress, and EMT in distal renal tubular epithelial cells [12, 60]. EMT is a critical pathological process in which epithelial cells lose their characteristic markers, acquire mesenchymal properties, and differentiate into myofibroblasts. These myofibroblasts actively contribute to renal fibrosis by producing extracellular matrix (ECM) components and fibrotic mediators, such as collagen and α-smooth muscle actin (α-SMA) [61, 62]. TGF-β1, a key mediator in the progression of CKD, plays a pivotal role in promoting fibrotic responses [62]. Additionally, CaOx crystals can trigger this fibrogenic cascade by upregulating markers, such as vimentin, α-SMA, and collagen, ultimately resulting in excessive ECM deposition and tubulointerstitial fibrosis [13, 15, 36]. Our data showed a significant increase in the expression of α-SMA, vimentin, and collagen-IV, along with a decrease in E-cadherin in both NRK-52E and PRTE cells exposed to COM exposure (**Fig. 9&10**). Similar to what we observed for other pathological changes after RIPK2 inactivation, a significant attenuation of α-SMA, vimentin, and collagen-IV expression following RIPK2/RICK silencing or inhibition suggests that RIPK2/RICK is a promising therapeutic target in the context of nephrocalcinosis associated EMT.

While our study provides valuable insights into the role of RIPK2/RICK in COM-induced renal injury, there are several limitations to consider. First, the use of *in vitro* model may not fully replicate the complexity of *in vivo* conditions, as the cellular responses observed *in vitro* may differ from those in a more dynamic *in vivo* environment. Additionally, while our findings are promising, the lack of *in vivo* validation limits the ability to confirm the therapeutic potential of targeting RIPK2/RICK in a whole organism context. Although our preliminary *in vivo* evidence does support RIPK2’s role in nephrocalcinosis, the *in vivo* pharmacological inhibition of RIPK2 is necessary to prove these findings. Our findings may apply to CaOx-associated nephrolithiasis, given their pathological similarities. Future studies utilizing *in vivo* COM models and advanced gene-editing techniques, such as CRISPR-Cas9, will address these concerns and provide a clearer understanding of RIPK2/RICK’s role in nephrocalcinosis and nephrolithiasis. This research has the potential to position RIPK2 as a therapeutic target for treating both nephrocalcinosis and nephrolithiasis.

## 5. Conclusion

Taken together, our study is the first to demonstrate that RIPK2/RICK expression is significantly upregulated in COM-challenged NRK-52E and PRTE cells. Silencing and inhibition of the RIPK2 notably attenuated COM-induced renal inflammation, oxidative stress, apoptosis, and EMT, likely through inhibition of the NF-κB/TGF-β signaling pathway. These findings highlight RIPK2/RICK as a promising therapeutic target for the prevention and treatment of nephrocalcinosis associated kidney damage.

## CRediT authorship contribution statement

**Ganesh Panditrao Lahane:** Writing – original draft, Methodology, Formal analysis. **Arti Dhar:** Writing – review & editing, Funding, Methodology. **Audesh Bhat:** Writing – review & editing, Funding, Methodology, Supervision, Conceptualization.

## Acknowledgements

We thank the Central Analytical Laboratory (CAL), BITS-Pilani Hyderabad Campus, for providing the necessary instrumentation facilities.

## Data Availability

All data supporting these findings are included within the manuscript.

---

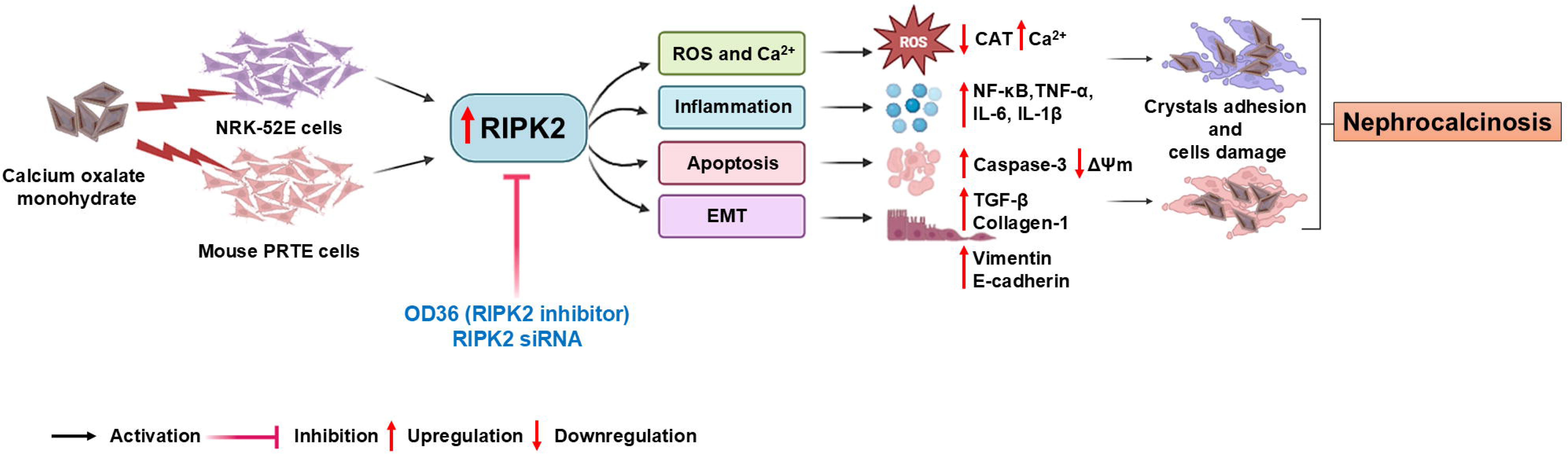

