## Supplementary material for "In-vitro Knockdown and Pharmacological Inhibition of RIPK2 Attenuates Calcium Oxalate–Nephrocalcinosis Associated Inflammation and Oxidative Stress in NRK-52E and Primary Renal Epithelial Cells": Suppl. Table 1

Table 1. Gene names and their primer sequences.

| **Species** | **Gene names** | **Forward primer (5′ −3′)** | **Reverse primer (5′ −3′)** |
| --- | --- | --- | --- |
| Mouse | CAT | GATGGTAACTGGGATCTTGTGG | GTGGGTTTCTCTTCTGGCTATG |
|  | IL-1β | GGTGTGTGACGTTCCCATTA | ATTGAGGTGGAGAGCTTTCAG |
|  | IL-6 | CTTCCATCCAGTTGCCTTCT | CTCCGACTTGTGAAGTGGTATAG |
|  | IL-10 | ACAGCCGGGAAGACAATAAC | CAGCTGGTCCTTTGTTTGAAAG |
|  | Tnf-α | CAGACCCTCACACTCACAAA | CTGGGAGTAGACAAGGTACAAC |
|  | Nf-κB | GGGATTTCGATTCCGCTATGT | GACCTGTGGGTAGGATTTCTTG |
|  | Caspase-3 | GACTGGAAAGCCGAAACTCT | GTATCTTCTGGCAAGCCATCT |
|  | Tgf-β | CTGAACCAAGGAGACGGAATAC | GGGCTGATCCCGTTGATTT |
|  | E-cadherin | GTCAAACGGCATCTAAAGC | CAAAGACCTCCTGGATAAACT |
|  | Vimentin | GATCGATGTGGACGTTTCCAA | GTTGGCAGCCTCAGAGAGGT |
|  | α-SMA | TCAGGGAGTAATGGTTGGAATG | GGTGATGATGCCGTGTTCTA |
|  | Collagen-IV (Col-IV) | TTACCTCCAACACCTCCAATC | GGGTCAAAGTACGGGTCATAG |
|  | β-actin | CAGCTGAGAGGGAAATCGTG | CGTTGCCAATAGTGATGACC |
|  | RIPK2/RICK | TTCCCTACCACAAACTCGCC | TCAGGCTCATTGCAAATTCCC |
| RAT | CAT | CATGGATCTGCTTAGGACTTCTG | CCAGGCTGTGAGGTAACATAA |
|  | IL-1β | GGAACCCGTGTCTTCCTAAAG | CTGACTTGGCAGAGGACAAA |
|  | IL-6 | GCTTTCGGAACTCACTGGAT | AGACATCTTCAGCAGCCTTG |
|  | IL-10 | CTGTCATTGATTTCTCCCCT | TCTATTTATGTCCTGCTGTCC |
|  | Tnf-α | GGAGAAGTTAGAGTCACAGAAGG | CACTAGGTTTGCCGAGTAGAC |
|  | Nf-κB | GGCTTCCTTTCTTGGCTCT | AAGCTCAAGCCACCATACC |
|  | Caspase-3 | TGGAAAGCATCCAGCAATAGG | GACTCAGCACCTCCATGATTAAG |
|  | Tgf-β | CTTTAGGAAGGACCTGGGTTG | GTGTCCAGGCTCCAAATGTA |
|  | E-cadherin | CCTCCTGCTCCTACTGTTTCTA | CTCCACCTCCCTCTTCATCATA |
|  | Vimentin | GCACGTCTTGACCTTGAACG | TGAGGTCAGGCTTGGAAACG |
|  | Α-SMA | AGGGAGTGATGGTTGGAATG | GGTGATGATGCCGTGTTCTA |
|  | Collagen-IV (Col-IV) | CACCCTGAACTCAAGAGCGG | TGCATGTTTCTCCGGTTTCC |
|  | β-actin | GAGGCCCCTCTGAACCCTAA | ACCAGAGGCATACAGGGAACAA |
